# Fallow deer approaching humans are also more likely to be seropositive for *Toxoplasma gondii*

**DOI:** 10.1101/2024.08.03.606274

**Authors:** Andrew R. Ryan, Annetta Zintl, Laura L. Griffin, Matthew Quinn, Amy Haigh, Pietro Sabbatini, Bawan Amin, Simone Ciuti

## Abstract

*Toxoplasma gondii* (*T. gondii*) is a trophically-transmitted protozoan parasite that has been suggested to facilitate its transmission by altering intermediate hosts’ anti-predator behaviour, thus increasing the likelihood of completing the cycle inside its definitive host i.e. domestic and wild felines. *T. gondii* has been linked to reduced risk-aversion, slower reaction times, and more exploratory behaviours in intermediate hosts, including most famously weakened aversion to the scent of feline predators in mice. Studies examining this phenomenon, however, have almost exclusively been carried out in laboratory conditions with small mammals, whereas little is known about the role of *T. gondii* within more complex ecological contexts involving large mammals in the wild. Under such scenario, the goals of our study were three-fold. Firstly, to determine the prevalence of *T. gondii* infection in a population of free-living fallow deer (*Dama dama*) living in a park at the edge of a metropolis. Secondly, to find a link between deer seropositivity and space use in the park, namely proximity to buildings with domestic cats, where deer may have been more likely to contract the disease. Finally, to determine whether infection with *T. gondii* was linked to risk taking behaviour in these free ranging large mammals, namely likelihood to approach park visitors. To achieve our goals, we estimated seropositivity and combined it with spatial distribution and behavioural data of individually-recognizable deer ranging from those that avoid humans (risk-avoiders) to those who beg for food (risk-takers). We found *T. gondii* to be quite widespread in this population with a seropositive of 20% (24 out of 120 individuals). Contrary to our expectations, we found no correlation between *T. gondii* seropositivity and space use in the park, therefore not allowing us to engage with the dynamics of disease contraction. We did however find that fallow deer taking the risk of approaching humans were also more likely to be seropositive. Are risk taking individuals more likely to contract the disease? Or, alternatively, do they take more risk because they have contracted the disease? The causal mechanism behind our result has yet to be disentangled, opening new scenarios in research aimed at tackling host manipulation in this parasite. It is a fact, however, that those animals that were more likely to be in contact with the public were also those more likely to be seropositive, adding key empirical evidence to the study of zoonotic diseases. Our study is a significant contribution on the transmission and maintenance dynamics of *T. gondii*, offering new insights on the need to conduct longitudinal studies able to disentangle the causal mechanism and *T. gondii*’s ability to manipulate its intermediate host.

## Introduction

It has been argued for some time that some trophically transmitted parasites (where the final host becomes infected by predation or scavenging on an intermediate host) have evolved the ability to manipulate their hosts’ behaviour to increase the likelihood to complete their life cycle (Lagrue and Poulin, 2010). One of the first parasites for which the host manipulation hypothesis was suggested was *Toxoplasma gondii* (Webster et al., 1994). *T. gondii* infects feline species as its only definitive hosts and most, if not all, warm-blooded animals as intermediate hosts (Elmore et al., 2010). Sexual reproduction in the epithelial cells of the feline small intestine results in the production of millions of oocysts which are released in the cat’s faeces (Torrey and Yolken, 2013). Under cool, moist conditions, oocysts can survive in the environment for months or even years (Frenkel et al., 1975). Once they are ingested by an intermediate host, the oocysts give rise to tachyzoites which invade various organs and tissues undergoing repeated rounds of asexual reproduction (Montoya and Liesenfeld, 2004). Eventually, the immune response of the intermediate host causes the tachyzoites to convert into metabolically less active bradyzoites which encyst in muscle and nervous tissues (Montoya and Liesenfeld, 2004). The cycle is completed if and when the intermediate host is consumed by another feline final host.

There are multiple methods by which *T. gondii* can be transmitted outside of this classical transmission cycle. For instance, non-feline predators or scavengers consuming intermediate hosts can be infected and act as another intermediate host (Lindsay and Dubey, 2020). *T. gondii* may also be transmitted vertically from mother to offspring. This however, may be host species-specific and has been recorded most commonly when the mother is infected during pregnancy. Vertical transmission has been documented in mice (Freyre et al., 2006), rats (Dubey and Shen, 1991), sheep (Williams et al., 2005 ), sea otters (Miller et al., 2008), and humans (Thiebaut et al., 2007).

It has been suggested that *T. gondii* may manipulate its intermediate hosts’ behaviour – namely increased risk-taking or a reduction in risk aversion - to make them more susceptible to predation, therefore increasing the likelihood that the final host eats the infected meat. Previous research has shown that the parasite may affect the synthesis of testosterone (Lim et al., 2013) and neurotransmitters such as dopamine (Prandovszky et al., 2011). Moreover, inflammatory responses to the presence of *T. gondii* cysts in brain tissue have also been associated with host behavioural changes (Tong et al., 2021). The best known examples of the host manipulation hypothesis reported in rodents demonstrated that *T. gondii* infections increased reaction times (Hrda et al., 2000), decreased neophobia (Webster et al., 1994), and reduced natural aversion to cat urine (Ingram et al., 2013). Tan and Vyas (2016) found that rats infected with *T. gondii* preformed closer to the optimal strategy when presented with balloon analogous risk task, due to a reduced level of risk aversion.

Comparable studies on free-living animals are less common, mostly due to difficulties in collecting epidemiological, diagnostic and behavioural data in wild settings simultaneously requiring a multi-disciplinary approach. While a range of publications have reported data on *T. gondii* prevalence in various species of wildlife (Lindsay and Dubey, 2020), fewer studies have examined potential behavioural changes. Gering et al. (2021) showed that wild hyena cubs infected with *T. gondii* come in closer proximity to lions which lead to higher mortality rates. Meyer et al. (2022) reported that wolves in Yellowstone National Park infected with *T. gondii* were more likely to take high risk decisions such as becoming a pack leader or dispersing from their pack. *T. gondii* has also been associated with more general central nervous system related pathologies, rather than the more specific behavioural changes mentioned above. Kreuder et al. (2003) found that Californian sea otters with encephalitis linked to *T. gondii* were 3.7 times more likely to be killed by shark attack. Similarly, *T. gondii* infections in wild red foxes have been associated with dopey fox syndrome (Milne et al., 2020).

As highlighted above, more studies are needed to disentangle the disease ecology and the role of *T. gondii* within more complex ecological contexts involving large mammals in the wild. To fill this gap, we conducted a study on a model population of fallow deer (*Dama dama*) living in one of the largest urban parks in Europe—the Phoenix Park in Dublin, Ireland. The goals of our study were three-fold. Firstly, to determine the seroprevalence of *T. gondii* infection in our model population, often interacting or spatially overlapping with humans and their pets because of the urban settings. Secondly, to determine whether there was a link between deer seropositivity and spatial proximity to buildings with domestic cats where deer would be more likely to contract the disease. Finally, to determine whether infection with *T. gondii* was linked to risk taking behaviour in these deer, namely the likelihood to approach park visitors. To achieve our goals we gathered unique data, collected adopting a multi-disciplinary approach involving behavioural and disease ecologists. We combined seropositivity with spatial distribution and behavioural data of individually-recognizable deer, ranging from those that avoid humans (risk-avoiders) to those who beg for food (risk-takers) over multiple years.

## Materials and methods

### Study site and population

The Phoenix Park is a 709-hectare walled park, situated within two kilometres of Dublin city centre. Two of the main habitats in the park are grasslands and assorted woodlands. The park hosts an estimated ten million visitors per year creating the ideal location to examine human-wildlife interactions (Griffin et al., 2022). The herd consists of approximately 600 fallow deer, of which circa 80% are identifiable via ear-tags that are applied each year to the majority of new-borns during the fawning season (Amin et al., 2021). As there are no predators capable of preying upon adult deer in the park, the population is managed by yearly culls.

Griffin et al. (2022) have documented in detail the behaviour of the majority of the individually recognizable deer of this population. In particular, focusing on their level of engagement with park visitors and willingness to tolerate human presence and close contact interactions to obtain processed human food (Griffin et al., 2022). Most of the deer usually avoid human contact and maintain a distance greater than 50 meters. A quarter of this population, however, tend to approach humans consistently and take advantage of additional artificial food sources (Griffin et al., 2022).

### Collection of blood samples

Blood samples were collected during the fawning period in June 2020 and June 2021 during the routine activities in the park when neonate deer are trapped and ear-tagged (Amin et al., 2021). Fallow deer adopt a “hider” strategy meaning that neonates hide in the vegetation while waiting for periodic visits of the mother for lactation, in general this behaviour lasts for at least 2-3 weeks after birth after which point the fawns generally join the herd (Chapman and Chapman, 1997). Precise data on the point at which fawns join the herd in this population can be found in Amin et al. (2022). For tagging, fawns are captured by a team of 10-15 people sweeping through the understory vegetation across the fawning sites of the park. When a fawn is spotted, it is caught using fishing nets, handled, and quickly released. Amin et al. (2021) includes a full description of the capture, handling and tagging operations following the highest standards of wildlife handling and welfare. As part of this process, small clips from the fawns’ ears were taken for genetic sampling, in most cases producing one or few drops of blood. These small amounts of blood were collected from the clip site using Nobuto blood filter-paper strip (Advantec 800700) as described by Nobuto (1966). The filter paper was then placed in individual sterile test tubes and kept on ice for transfer into a freezer by the end of the day. All animal handling was conducted with permission from the UCD Animal Research Ethics Committee, under the permit AREC-E-18-28. The serostatus of the fawns was used to indirectly determine whether their mothers had antibodies against *T. gondii*. The identity of the mothers was determined via behavioural observations (e.g., suckling, grooming, following between the mother and the fawn) as described in Griffin et al. (2022) and Amin et al. (2022).

In addition blood samples were collected during the annual culls of 2020 and 2021, in the period between November and January. The cull is conducted by professional stalkers hired by the government body responsible for the management of deer in the park, the Office of Public Work (OPW), under a hunting permit issued by the National Park and Wildlife Services (NPWS). During the cull, we collected blood samples in 50ml falcon tube as the carcass was hung up. Sera were collected following centrifugation (2500g for 10 minutes) and stored at -20°C. In contrast to the fawning blood sample collection, where we indirectly sampled mothers corresponding to the fawns (thus, only females were indirectly sampled), we directly collected both male and female deer sera samples during the cull. The cull targeted individual deer at random following a precise culling plan with quotas indicating the minimum number of age and sex classes to be removed from the population.

### ELISA test: seropositivity in the sampled deer population

Sera were tested for the presence of specific antibody using the ID Screen® Toxoplasmosis Indirect Multi-Species Enzyme-Linked Immunosorbent Assay (ELISA) kit (Product code: TOXOS-MS-2P). For the fawn samples collected on Nobuto blood filter-paper strips, sera were eluted from filter paper squares by soaking them overnight in the dilution buffer provided by the kit. Depending on the amount of blood that had been captured on the filter paper, samples consisted of 0.1, 0.5, 1, or 2 squares of dimensions 0.5cm x 0.5cm. Samples with two squares were eluted in 200 µL of dilution buffer, samples with one square were eluted in 150 µL of dilution buffer, samples with 0.5 squares were eluted in 100µL of dilution buffer, respectively. All samples were applied directly to the test plate (i.e. analysed without further dilution) and each sample was in duplicate.

The serum samples derived from the blood collected during the deer cull were analysed at a dilution of 1/10 as recommended by the kit manufacturer. Again, two replicates were tested for each sample. Positive and negative controls provided in the kit were included on each plate. For each sample, a Sample / Positive percentage (S/P%) was calculated from the optical density (OD) values using the following formula:

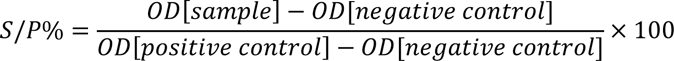

According to the kit manufactures, S/P% values less than 40 are considered negative, S/P% values from 40 to 50 are equivocal, and S/P% values greater than 50 are positive. In order to determine whether slight differences in the blood sample on the filter paper and the volume of elution buffer would affect the outcome of the analysis, a panel of 11 deer sera were analysed both at dilutions of 1/2 and 1/10. While S/P% values increased somewhat at the lower dilution factor, the test outcome was mostly unaffected (results in Supplementary material S1).

### Analysis of space use by deer in the park with respect to buildings

The space use of individual deer was determined using a survey carried out on a weekly or bi-weekly basis from September 2018 to December 2021 by tracking the spatial location of individually-recognizable deer using a hand-held GPS unit combined with the use of a compass and a rangefinder. For much of the year males and females use different sections of the park (Griffin et al., 2022) so they were examined separately. Data on seropositivity (either positive or negative to *T. gondii*) were combined with spatial data by generating heatmaps using the *Leaflet* (Cheng et al., 2022) and *Leaflet.extras* (Karambelkar and Schloerke, 2018) packages in R (version 4.1.3) (R Core Team, 2022). Firstly, a qualitative approach was used to determine whether the distribution of seropositive vs seronegative deer differed on a large scale (i.e., whether positive individuals were clearly using different areas compared to negative individuals). Secondly, in a more quantitative approach, we fitted a linear mixed-effect model (*lmer* function of the *lme4* package, (Bates et al., 2015)) to test whether seropositive individuals spent more time closer to buildings in the park, where they were more likely to become exposed to oocysts deriving from cat faeces. An accurate map of buildings in the park was obtained from Ordinance Survey Ireland (OSI, the Irish state’s national mapping body). We fitted an a priori mixed effect model using the following formula:

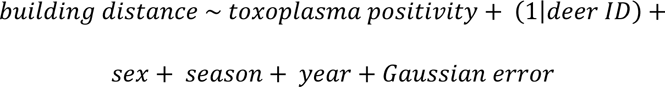

where: building distance is a numeric value of the distance in meters from the nearest building for each GPS location where individual deer were recorded during the regular park surveys; toxoplasma seropositivity is a categorical variable indicating whether animals tested “Positive” or “Negative”; deer ID is the random intercept denoting the individual identification tag of each individual deer; sex is a categorical variable that distinguishes between “Males” and “Females”; season is a categorical variable indicating the time of the year as the vast majority of males join the female heard during the rut. Season is described as “Winter/Spring” (January to April), “Summer” (May to August), and “Rut” (September to December). Finally, year is a categorical variable that differentiates between the different years of location data collection (2018, 2019, 2020, 2021). Year is included to account for any differences in the environment (such as vegetation cover or visitor numbers) that could have occurred in the park during the study period.

### Begging ranks: tendency of deer to beg for food sensu Griffin et al. (2022)

We obtained the begging ranks (on a continuum from deer avoiding any interaction with humans to those persistently seeking contact and begging for food) based on field observations taken over the summers of 2020 and 2021, in the period immediately preceding and simultaneous to our blood data collection. Full data collection protocol and data analysis can be found in Griffin et al. (2022). In brief, deer in the park were systematically surveyed and documented avoiding humans or taking food from them. The best linear unbiased predictors (BLUPs, i.e. the random effect of a Generalized Linear Mixed Effect model explaining the variability of begging behaviour) were extracted from Griffin et al. (2022)’s analysis and used as begging rank depicting the willingness of deer to approach humans and accept related risks. Begging rank (mean: 0.58; 1^st^ quantile: -0.23; 3^rd^ quantile: 1.49) varied from -2.40 (min value, i.e., the individual avoiding humans the most) to 4.57 (max value, i.e., the individual in the sample population begging for food the most).

### Modelling begging rank as a function of seropositivity

Each ELISA reading, expressed as S/P%, was used as a data point. To this value we attached the rest of the data such as the individual deer ID, sex, year of collection, begging rank of the individual for that year, and the begging rank uncertainty *sensu* Griffin et al. (2022). The begging rank uncertainty, in particular, is an important confounding factor which estimates how robust the estimate of the begging rank is based on sample size and number of observations per individual deer (Griffin et al., 2022). We fitted a liner mixed-effect model to explain the variability of begging rank (response variable representing the willingness of deer to take the risk and approach park visitors) as a function of the S/P% values, our main predictor, and a set of confounding covariates: sex, begging rank uncertainty, replicate number, year of data collection. Type of sample (i.e. filter paper-eluted samples obtained from fawns vs sera collected from culled deer) was omitted from the model equation due to its collinearity with sex (Dormann et al., 2013).

We log-transformed our S/P% values to improve the model’s fit and to better meet the model’s assumptions of residual normality and homogeneity and we scaled all numeric predictors (i.e., by subtracting the mean and dividing by the standard deviation) to improve the model’s convergence. Our model equation was:

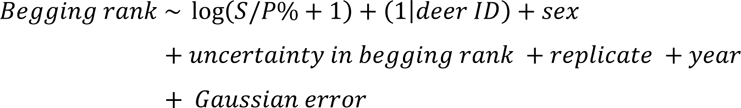

where: deer ID was fitted as a random intercept in the model, S/P% and begging rank uncertainty as numerical predictors, and replicate and year of study were fitted as categorical predictors. Linear regression models were created using the *lme4* package (Bates et al., 2015) in R (version 4.1.3)(R Core Team, 2022).

## Results

### T. gondii seropositivity

Over two fawning seasons we obtained samples from 139 fawns that were tested by ELISA. This was split into 53 fawns in 2020 and 86 fawns in 2021 (seropositivity results for both years in Table 1). As the fawns will have obtained the antibodies they are expressing from their mothers and fallow deer in the Phoenix park have never been recorded having more than one offspring per year (unpublished data) we can be confident in reporting the seropositivity rates of all fawns in a single year without including multiple results from the same individual. However, when combining results from both years it is important that we can identify the mother of each fawn to attribute the results to her. From the fawns we tested we were able to confidently identify 64 mothers. Seventeen mothers were tested in both fawning seasons with 4 being removed as their serostatus changed during the study (details below). Of the remaining 60 does; 18 (30%) tested positive, 29 (48.3%) tested negative, and 13 (21.7%) gave equivocal results. The age of the mothers tested through their fawns spread from a minimum of 2 years to a maximum of 16 years with an average value of 6.4 years.

**Table 1:** Serological screen of blood samples collected from fawns during ear notching and from adult deer during the annual cull. This includes both tagged and untagged individuals unless otherwise stated. Results are based on replicate samples tested using ELISA. Samples with an S/P%>50% were considered positive and samples with S/P%<40% were considered negative. Samples with S/P% from 40% to 50% or where the two replicates did not match were considered equivocal. Seroprevalence in does during fawning was estimated by matching individuals fawns with their mothers.

| Sampling occasion | Year of sample collection | N | Positive | Negative | Equivocal |
| --- | --- | --- | --- | --- | --- |
| Fawning | 2020 | 53 | 13 (24.5%) | 34 (64.2%) | 6 (11.3%) |
| Fawning | 2021 | 86 | 24 (27.9%) | 33 (38.4%) | 29 (33.7%) |
| Combined Fawning (Identifiable does only) | 2020 & 2021 | 60 | 18 (30%) | 29 (48.3%) | 13 (21.7%) |
| Cull | 2020 | 46 | 5 (10.9%) | 37 (80.4%) | 4 (8.7%) |
| Cull | 2021 | 42 | 7 (16.7%) | 31 (73.8%) | 4 (9.5%) |
| Combined Cull | 2020 & 2021 | 88 | 12 (13.6%) | 68 (77.3%) | 8 (9.1%) |

Out of the 88 deer sampled during the annual culls; 12 (13.6%) tested positive, 68 (77.3%) tested negative and 8 (9.1%) were equivocal. Seroprevalence of unequivocal samples were similar between the sexes with 12.8% seropositivity in males (5 out of 39 samples) and 17.5% seropositivity in females (7 out of 40 samples) for one animal the sex was not recorded. Nineteen of the culled deer were excluded from further analysis as they had no ear tag, therefore we could not recover any life history information for them regarding begging behaviour or movements within the park. This left us with 61 individuals, 8 of which tested positive (13.1%) with ages ranging from 0 (fawns born approximately six months before the cull) to 11 years with an average age of 2.6 years.

Overall 227 samples tested by ELISA during this study. Seventy eight of these samples derived from untagged deer, as we cannot track which individual deer these came from we cannot be sure if deer were tested multiple times. The remaining 149 samples came from 125 tagged individuals. Of these tagged individuals 102 were sampled in one sampling window (21 testing positive, 66 negative, and 15 equivocal). Twenty-two tagged deer were sampled during two sampling windows; 9 tested negative both times, 5 tested negative once and equivocal once, 3 tested positive once and equivocal once, 4 tested negative once and positive once, and 1 tested equivocal both times. One individual (Yellow A26) was tested in three sampling windows; twice indirectly - she had a fawn that tested negative in June 2020 and a fawn that tested positive in June 2021- and then again during the cull in November 2021 where she tested negative. All animals that changed their serostatus during the course of the study were excluded further analysis as it was not possible to determine when exactly they became exposed (and turned seropositive) or cleared the parasite (i.e. turned seronegative). Our overall estimate for seropositivity in this population, based solely on tagged deer, is 20% (24 positive, 80 negative (66.6%), and 16 equivocal (13.3%) out of 120 individuals).

### Association between spatial behaviour and T. gondii serostatus

Figure 1 shows heatmaps created to visualize space use seronegative (n= 48) (A) and seropositive (n= 21) (B) female deer. There was as strong overlap in space use between the two groups and there were no areas predominantly used by seropositive individuals that were not used by seronegative deer and vice versa. Similar results were obtained for the male deer (results shown in Supplementary material: Males; S2 A&B, Females; S2 C&D).

**Fig. 1:**
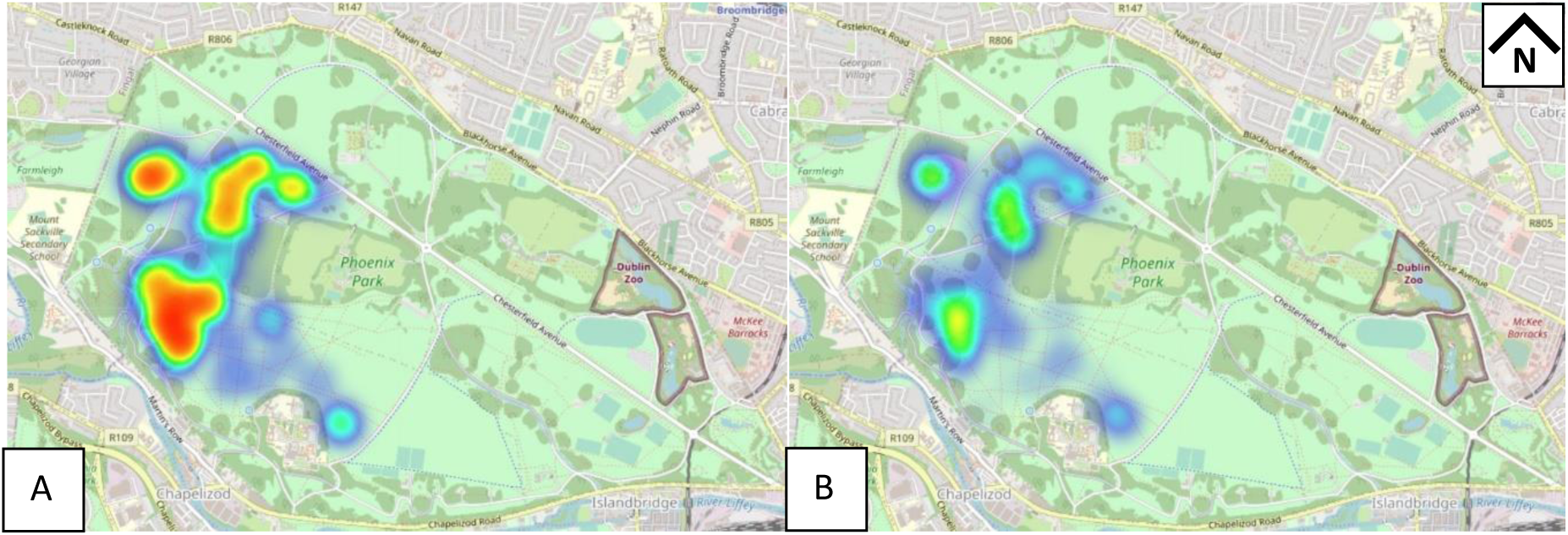
Heatmaps showing space by (A) seronegative and (B) seropositive female deer in Phoenix Park, Dublin. The male heatmaps are included in the supplementary materials.

Table 2 provides the results of the mixed effect model explaining the variability of the distance to the buildings. There was no clear link between seropositivity and distance from buildings (see Supplementary material S3 for model effect plots). Males were generally found closer to buildings than females (Table 2, S3), whereas both sexes were found to be closer to buildings during the rut (Table2, S3) than during the rest of the year.

**Table 2:** Parameters estimated by the linear mixed effect model explaining the distance from buildings of a deer observed at a point as a function of *T. gondii* status and a range of confounding variables, including sex, season and year. Sex distinguishes between males and females. Serostatus denotes seropositive and seronegative individuals. Season was described as Winter/Spring (January to April), Summer (May to August), and Rut (September to December). Year included the four years of observations: 2018, 2019, 2020, and 2021.

| Predictors | Estimates | Confidence |  |
| --- | --- | --- | --- |
|  |  | Interval | p-value |
| Intercept | 228.24 | 195.15 – 261.34 | **<0.001** |
| Sex [Male] | -38.50 | -62.71 – -14.29 | **0.002** |
| Serostatus [Positive] | -3.78 | -29.48 – 21.92 | 0.773 |
| Season [Summer] | 46.57 | 19.48 – 73.67 | **0.001** |
| Season [Winter/Spring] | 45.37 | 20.66 – 70.09 | **<0.001** |
| Year [2019] | -11.73 | -49.79 – 26.34 | 0.546 |
| Year [2020] | -6.85 | -46.00 – 32.30 | 0.731 |
| Year [2021] | -35.44 | -76.60 – 5.72 | 0.091 |
| Random Effects |  |  |  |
| $\sigma^2$ | 17906.26 | | |
| $\tau^2_{00\text{ ID}}$ | 933.20 | | |
| ICC | 0.05 |  |  |
| $N_{ID}$ | 103 | | |
| Observations | 856 |  |  |
| Marginal $R^2$/Conditional $R^2$ | 0.036/0.084 | | |

### Begging rank as a function of S/P% ratios

Results from the mixed-effect model explaining the variability of begging rank as a function of ELISA S/P% and a range of confounding factors are shown in Table 3. The analysis was performed including each ELISA S/P% result (i.e. including both replicates) rather than the average of the two (thus including a two-level predictor “replicate” as controlling factor) . Individuals with higher S/P% values were associated with an increased tendency to beg for food (Table 3, Fig. 2). Females were more likely to be found seropositive (Fig. 3a) after accounting for begging rank uncertainty and sample replicate (Fig. 3b,c), and in 2021 we found a greater level of seropositivity than the previous year (Fig. 3d).

**Fig. 2:**
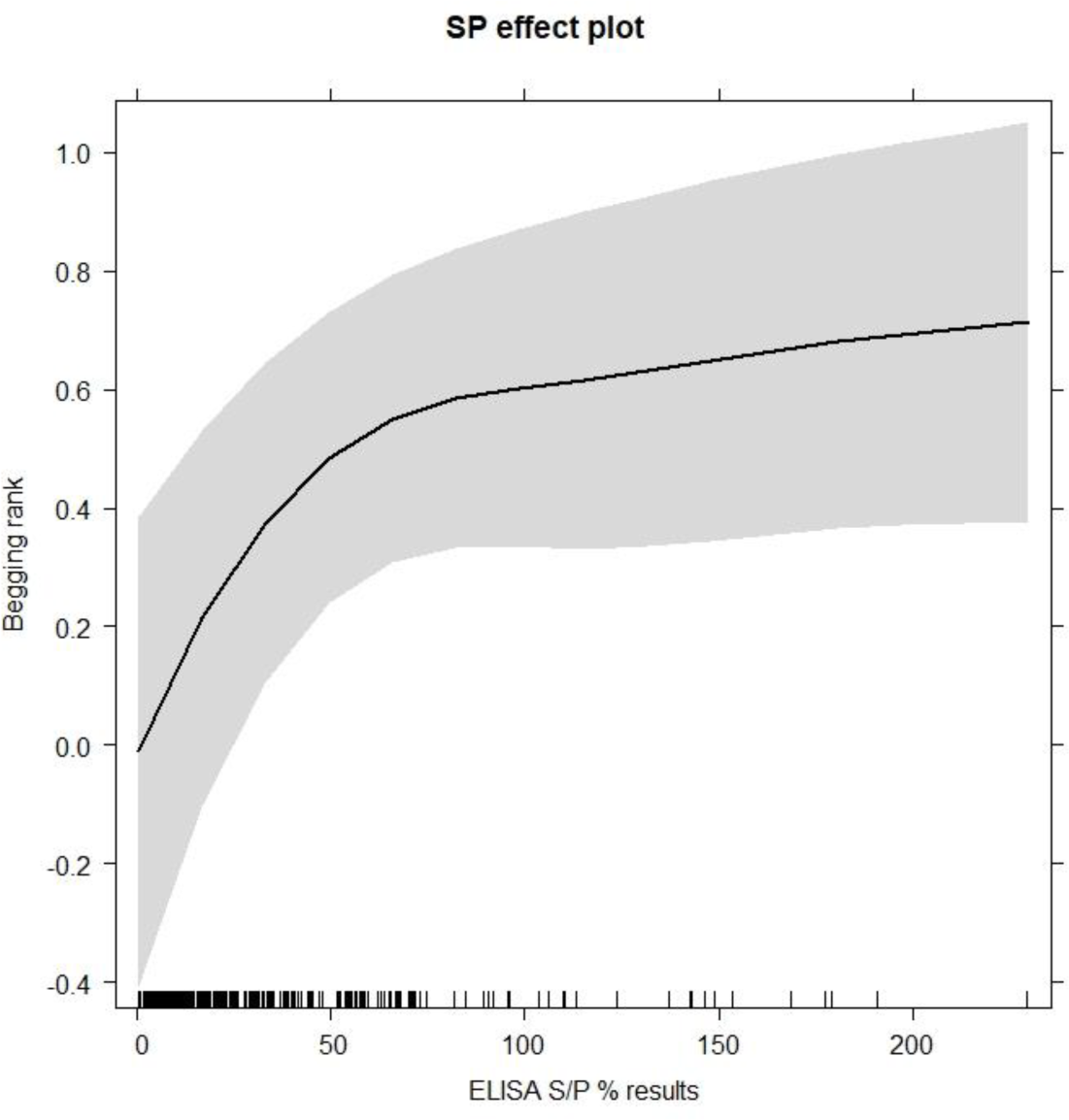
Effect plot depicting the relationship between ELISA S/P% (x-axis) and begging rank (y-axis, with greater ranks corresponding to higher likelihood to interact and accept food from park visitors) as predicted by the linear mixed effect model. The shaded area represents the 95% marginal confidence intervals.

**Fig. 3:**
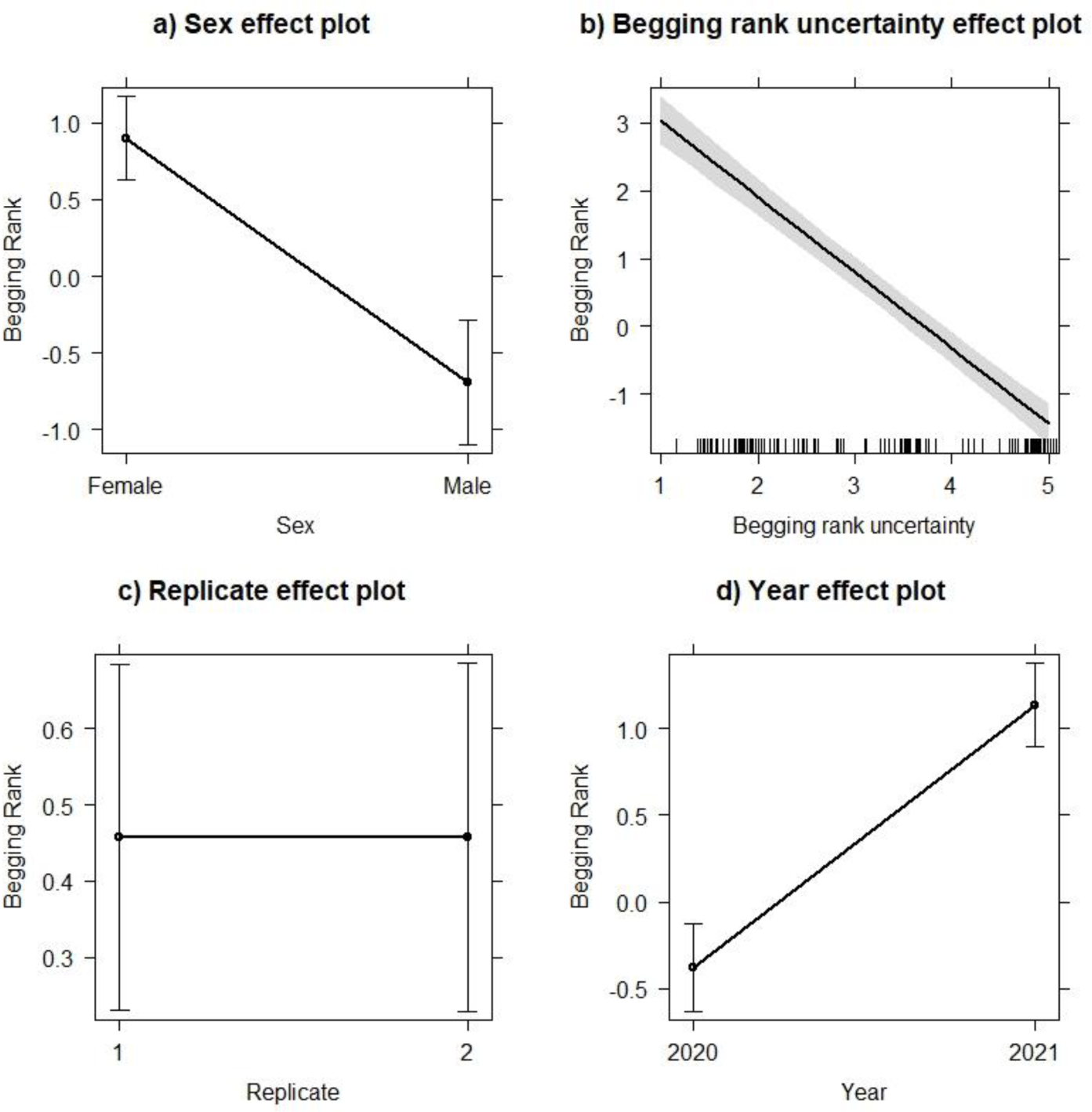
Effect plot depicting the effect of sex (a), begging rank uncertainty (b), replicate test (c), and year of study (d) on tendency to beg (y-axes) in deer monitored in the Phoenix Park, Dublin, as predicted by a linear mixed-effect model. The bars / shaded area represents the 95% marginal confidence intervals.

**Table 3:** Parameters estimated by the linear mixed effect model explaining the variability of begging rank in deer as a function of ELISA S/P% and a range of confounding variables. The begging rank is a score associated to individual deer based on their likelihood to beg, with higher values meaning more likely to beg. S/P% are the numeric values obtained from the ELISA test results. Sex is a factor which distinguishes between male and female individuals. Begging rank uncertainty (i.e. the uncertainty arising from the begging rank estimates) ranges from 1.2 to 5.1. Each sample was tested in duplicate and the analysis was performed to include each ELISA S/P% result rather than the average, the factor replicate distinguished between the first and second replicates. The study was run with samples from two years (2021 and 2022).

| Predictors | Estimates | Confidence Interval | p-value |
| --- | --- | --- | --- |
| Intercept | 0.01 | -0.27 – 0.29 | 0.945 |
| S/P% | 0.13 | 0.02 – 0.23 | **0.016** |
| Sex [Male] | -1.59 | -2.08 – -1.11 | **<0.001** |
| Begging rank uncertainty | -1.37 | -1.52 – -1.23 | **<0.001** |
| Replicate [run 2] | 0.00 | -0.07 – 0.07 | 0.992 |
| Year [2021] | 1.51 | 1.31 – 1.71 | **<0.001** |
| Random Effects |  |  |  |
| $\sigma^2$ | 0.08 | | |
| $\tau^2_{00}$ ID | 1.37 | | |
| ICC | 0.94 |  |  |
| $N_{ID}$ | 115 | | |
| Observations | 259 |  |  |
| Marginal $R^2$/Conditional $R^2$ | 0.474/0.970 | | |

## Discussion

Seroprevalence of *T. gondii* in the fallow deer population in the Phoenix Park differed significantly depending on the timing and cohort of sampling. Samples collected from fawns, which were used to estimate the serostatus of their tagged mothers at the time of fawning, had a seropositivity rate of 30%. As deer are ruminants, mothers pass antibodies to their fawns through their colostrum only during the first few feeds, not across the placenta (Butler and Kehrli, 2005). Therefore, testing the blood samples of these fawns (which are usually less than 1 week old), can serve as an indirect method to determine whether the mothers are seropositive for *T. gondii*. In contrast, sera from culled deer had a seropositivity rate of just 13.6%. The same difference was observed at repeated sampling occasions (seroprevalence at fawning in 2020 was 24.5% and 27.9% in 2021; seroprevalence at the cull was 10.9% in 2020 and 16.7% in 2021). The age of the mothers of the fawns that were tested was considerably higher (range of 2 to 16 years, average of 6.4 years) than that of the culled deer (range of 0 to 11 years, average: 2.6 years) and older animals are generally more likely to be seropositive for *T. gondii* as they have had more opportunity to become exposed over time (Wilking et al., 2016, Katzer et al., 2011, Chadwick et al., 2013). Another likely explanation (possibly additive to the effect of age) is that periparturient immunosuppression in pregnant does may allow dormant *T. gondii* bradyzoites resident tissue cysts to reactivate sufficiently to cause an increase in serum antibody levels. Reactivation of chronic infections during pregnancy have been documented in humans (Biedermann et al., 1995) and suspected in sheep and goats (Sánchez-Sánchez et al., 2018, Williams et al., 2005 ). Moreover, periparturient immune suppression and an associated increase in the shedding of certain parasites has previously been reported for other deer species: red deer (Skerrett and Holland, 2001) and sika deer (Sarkunas et al., 2007).

We provided empirical evidence on the fact that the population of fallow deer in Phoenix Park has several *T. gondii* seropositive individuals. The most likely method for herbivorous animals to be infected is through ingestion of oocysts from the faeces of infected cats. In fact it has been reported that rodents caught closer to farm buildings where there were more cats, and consequently more oocysts, are more likely to be infected with *T. gondii* (Gotteland et al., 2014a, Gotteland et al., 2014b). Therefore, before tackling our final and most important question on the association between *T. gondii* serostatus and risk taking behaviour, we first had to determine whether animals that were naturally less risk averse were more likely to spend time near buildings and become exposed to oocysts. Contrary to our expectations, there was no correlation between serostatus and distance from buildings. Both qualitative and quantitative analyses showed seropositive and seronegative animals sharing the same areas. We therefore do not have empirical evidence on how deer in this population have been exposed to the parasite, and we can only discuss a number of speculative theories that would need further research, both in peri-urban and rural deer populations. Besides buildings and associated domestic cats, there are two possible pathways for deer to be exposed to *T. gondii* which may be through infectious material brought by visitors, i.e. (i) raw meat containing tissue cysts and/or (ii) shoes/clothes contaminated with cat faeces. The risk can be considered very low, but cannot be ruled out. In Ireland relatively low levels of *T. gondii* infections have been reported in livestock (Halova et al., 2013). Through previous research in the Phoenix Park there are scattered records of the deer accepting cooked meat included in sandwiches offered by park visitors (Griffin et al., 2022); however, there have been no observations of visitors feeding deer uncooked meat. As *T. gondii* cannot survive in cooked meat, it would be improbable that any meat fed to the deer would be both uncooked and infected with *T. gondii*. On the other hand, it is quite unlikely (but not impossible) that the number of viable oocysts transported via shoe/hand contamination would suffice to infect an immune-competent deer. More research is required, as we could find no reports in the literature examining the transport of oocysts by humans on their hands, clothing, or shoes.

The main aim of this study was to verify whether observations of potential host manipulation by *T. gondii* reported rodents under laboratory conditions (Webster et al., 1994, Berdoy et al., 2000, Ingram et al., 2013) translate to free living, non-rodent hosts. In the Phoenix Park, humans are the only major potential threat faced regularly by the deer through annual culls aimed at maintaining a healthy and sustainable population withing a walled park. Deer are generally habituated to human presence but avoid any close contact, with the exception of a few individuals that get closer to humans (Griffin et al., 2022). Our analysis showed a significant association between *T. gondii* serostatus and begging behaviour, i.e. the tendency of some individuals to win the fear of humans and get closer to take advantage of artificial food items. *T. gondii* antibody levels were indeed positively correlated with begging behaviour indicating that individuals that had been exposed to *T. gondii* were bolder (less risk adverse) than their unexposed counterparts, in line with our main hypothesis on host behavioural manipulation.

Despite this very interesting result, we refrain ourselves to boldly claim that the behaviour of fallow deer is manipulated by *T. gondii*. Our result is novel, adds important empirical evidence on the dynamics of zoonotic disease maintenance and transmission, but we are not in the position to draw final conclusions about the causal relationship of our result. On the one hand, the likelihood to adopt risk taking behaviours and approach humans may be boosted by *T. gondii* infection, along with individual temperament or personality (Amin et al., 2021, Griffin et al., 2022). On the other hand, it may be possible that those risk-taking bolder individuals (*sensu* Griffin et al. (2022)) were more likely to contract the disease from humans. Despite the latter scenario seems unlikely, it cannot be excluded with certainty. Only long-term longitudinal behavioural studies, where deer monitored for years would start engaging in risk-taking behaviour only after contracting the parasite will allow us to draw final conclusions on the hypothesis on *T. gondii* manipulating its host. From a zoonotic disease perspective, our study demonstrates that park visitors are more likely to interact with seropositive individuals. Likewise, hunters may be more likely to shoot these bolder individuals as previously shown by Ciuti et al. (2012). Hunters often handle raw meat without any sort of protection, therefore with a possibility for *T. gondii* to pass from an intermediate host to another. Toxoplasmosis has been shown to be widespread across deer populations (see Keenan et al. (2024)), and our study highlights that humans are likely to interact with seropositive individuals, suggesting that biosecurity measures should be a practice particularly when handling the raw meat of these animals.

## References

Amin, B., Jennings, D. J., Norman, A., Ryan, A., Ioannidis, V., Magee, A., Haughey, H.-A., Haigh, A. & Ciuti, S. 2022. Neonate personality affects early-life resource acquisition in a large social mammal. Behavioral Ecology, 33, 1025–1035. doi:10.1093/beheco/arac072

Amin, B., Jennings, D. J., Smith, A. F., Quinn, M., Chari, S., Haigh, A., Matas, D., Koren, L. & Ciuti, S. 2021. In Utero Accumulated Steroids Predict Neonate Anti-Predator Response in a Wild Mammal. Functional Ecology, 35, 1255–1267. doi:10.1111/1365-2435.13790

Bates, D., Mächler, M., Bolker, B. M. & Walker, S. C. 2015. Fitting Linear Mixed-Effects Models Using lme4. Journal of Statistical Software, 67, 1–48. doi:10.18637/jss.v067.i01

Berdoy, M., Webster, J. P. & Macdonald, D. W. 2000. Fatal attraction in rats infected with *Toxoplasma gondii*. Proceedings of the Royal Society B-Biological Sciences, 267, 1591–1594. doi:10.1098/rspb.2000.1182

Biedermann, K., Flepp, M., Fierz, W., Jollerjemelka, H. & Kleihues, P. 1995. Pregnancy, immunosuppression and reactivation of latent toxoplasmosis. Journal of Perinatal Medicine, 23, 191–203. doi:10.1515/jpme.1995.23.3.191

Butler, J. E. & Kehrli, M. E. 2005. Immunoglobulins and Immunocytes in the Mammary Gland and Its Secretions. Mucosal Immunology, *3rd Edition.* doi:10.1016/b978-012491543-5/50107-8

Chadwick, E. A., Cable, J., Chinchen, A., Francis, J., Guy, E., Kean, E. F., Paul, S. C., Perkins, S. E., Sherrard-Smith, E., Wilkinson, C. & Forman, D. W. 2013. Seroprevalence of *Toxoplasma gondii* in the Eurasian otter (*Lutra lutra*) in England and Wales. Parasites & Vectors, 6. doi:10.1186/1756-3305-6-75

Chapman, D. & Chapman, N. 1997. *Fallow Deer: Their History, Distribution and Biology*, Coch-y-Bonddu Books. isbn:9780952851059.

Cheng, J., Karambelkar, B. & Xie, Y. 2022. leaflet: Create Interactive Web Maps with the JavaScript ’Leaflet’ Library, R package version 2.1.1. https://CRAN.R-project.org/package=leaflet.

Ciuti, S., Muhly, T. B., Paton, D. G., Mcdevitt, A. D., Musiani, M. & Boyce, M. S. 2012. Human selection of elk behavioural traits in a landscape of fear. Proceedings of the Royal Society B-Biological Sciences, 279, 4407–4416. doi:10.1098/rspb.2012.1483

Dormann, C. F., Elith, J., Bacher, S., Buchmann, C., Carl, G., Carre, G., Marquez, J. R. G., Gruber, B., Lafourcade, B., Leitao, P. J., Munkemuller, T., Mcclean, C., Osborne, P. E., Reineking, B., Schroder, B., Skidmore, A. K., Zurell, D. & Lautenbach, S. 2013. Collinearity: a review of methods to deal with it and a simulation study evaluating their performance. Ecography, 36, 27–46. doi:10.1111/j.1600-0587.2012.07348.x

Dubey, J. P. & Shen, S. K. 1991. Rat model of congenital toxoplasmosis. Infection and Immunity, 59, 3301–3302. doi:10.1128/iai.59.9.3301-3302.1991

Elmore, S. A., Jones, J. L., Conrad, P. A., Patton, S., Lindsay, D. S. & Dubey, J. P. 2010. *Toxoplasma gondii*: epidemiology, feline clinical aspects, and prevention. Trends in Parasitology, 26, 190–196. doi:10.1016/j.pt.2010.01.009

Frenkel, J. K., Ruiz, A. & Chinchilla, M. 1975. Soil survival of toxoplasma oocysts in Kansas and Costa-Rica. American Journal of Tropical Medicine and Hygiene, 24, 439–443. doi:10.4269/ajtmh.1975.24.439

Freyre, A., Falcon, J., Mendez, J., Rodriguez, A., Correa, L. & Gonzalez, M. 2006. Refinement of the mouse model of congenital toxoplasmosis. Experimental Parasitology, 113, 154–160. doi:10.1016/j.exppara.2005.12.019

Gering, E., Laubach, Z. M., Weber, P. S. D., Hussey, G. S., Lehmann, K. D. S., Montgomery, T. M., Turner, J. W., Perng, W., Pioon, M. O., Holekamp, K. E. & Getty, T. 2021. *Toxoplasma gondii* infections are associated with costly boldness toward felids in a wild host. Nature Communications, 12. doi:10.1038/s41467-021-24092-x

Gotteland, C., Chaval, Y., Villena, I., Galan, M., Geers, R., Aubert, D., Poulle, M. L., Charbonnel, N. & Gilot-Fromont, E. 2014a. Species or local environment, what determines the infection of rodents by *Toxoplasma gondii*? Parasitology, 141, 259–268. doi:10.1017/s0031182013001522

Gotteland, C., Mcferrin, B. M., Zhao, X. P., Gilot-Fromont, E. & Lelu, M. 2014b. Agricultural landscape and spatial distribution of *Toxoplasma gondii* in rural environment: an agent-based model. International Journal of Health Geographics, 13. doi:10.1186/1476-072x-13-45

Griffin, L. L., Haigh, A., Amin, B., Faull, J., Norman, A. & Ciuti, S. 2022. Artificial selection in human-wildlife feeding interactions. Journal of Animal Ecology, 91, 1892–1905. doi:10.1111/1365-2656.13771

Halova, D., Mulcahy, G., Rafter, P., Turcekova, L., Grant, T. & De Waal, T. 2013. *Toxoplasma gondii* in Ireland: Seroprevalence and Novel Molecular Detection Method in Sheep, Pigs, Deer and Chickens. Zoonoses and Public Health, 60, 168–173. doi:10.1111/j.1863-2378.2012.01514.x

Hrda, S., Votypka, J., Kodym, P. & Flegr, J. 2000. Transient nature of *Toxoplasma gondii*-induced behavioral changes in mice. Journal of Parasitology, 86, 657–663. doi:10.1645/0022-3395(2000)086[0657:tnotgi]2.0.co;2

Ingram, W. M., Goodrich, L. M., Robey, E. A. & Eisen, M. B. 2013. Mice Infected with Low-Virulence Strains of *Toxoplasma gondii* Lose Their Innate Aversion to Cat Urine, Even after Extensive Parasite Clearance. Plos One, 8. doi:10.1371/journal.pone.0075246

Karambelkar, B. & Schloerke, B. 2018. leaflet.extras: Extra Functionality for ’leaflet’ Package. R package version 1.0.0. https://CRAN.R-project.org/package=leaflet.extras.

Katzer, F., Brulisauer, F., Collantes-Fernandez, E., Bartley, P. M., Burrells, A., Gunn, G., Maley, S. W., Cousens, C. & Innes, E. A. 2011. Increased *Toxoplasma gondii* positivity relative to age in 125 Scottish sheep flocks; evidence of frequent acquired infection. Veterinary Research, 42. doi:10.1186/1297-9716-42-121

Keenan, S., Niedziela, D., Morera-Pujol, V., Franklin, D., Murphy, K. J., Ciuti, S. & Mcmahon, B. J. 2024. Zoonotic disease classification in wildlife: a theoretical framework for researchers. Mammal Review, 54, 63–77. doi:10.1111/mam.12329

Kreuder, C., Miller, M. A., Jessup, D. A., Lowenstein, L. J., Harris, M. D., Ames, J. A., Carpenter, T. E., Conrad, P. A. & Mazet, J. A. K. 2003. Patterns of mortality in southern sea otters (*Enhydra lutris nereis*) from 1998-2001. Journal of Wildlife Diseases, 39, 495–509. doi:10.7589/0090-3558-39.3.495

Lagrue, C. & Poulin, R. 2010. Manipulative parasites in the world of veterinary science: Implications for epidemiology and pathology. Veterinary Journal, 184, 9–13. doi:10.1016/j.tvjl.2009.01.015

Lim, A., Kumar, V., Dass, S. A. H. & Vyas, A. 2013. *Toxoplasma gondii* infection enhances testicular steroidogenesis in rats. Molecular Ecology, 22, 102–110. doi:10.1111/mec.12042

Lindsay, D. S. & Dubey, J. P. 2020. Toxoplasmosis in wild and domestic animals. *In:* Weiss, L., M. & Kim, K. (eds.) Toxoplasma gondii: the Model Apicomplexan-Perspectives and Methods, *3rd Edition.* doi:10.1016/b978-0-12-815041-2.00006-2

Meyer, C. J., Cassidy, K. A., Stahler, E. E., Brandell, E. E., Anton, C. B., Stahler, D. R. & Smith, D. W. 2022. Parasitic infection increases risk-taking in a social, intermediate host carnivore. Communications Biology, 5. doi:10.1038/s42003-022-04122-0

Miller, M., Conrad, P., James, E. R., Packham, A., Toy-Choutka, S., Murray, M. J., Jessup, D. & Grigg, M. 2008. Transplacental toxoplasmosis in a wild southern sea otter (*Enhydra lutris nereis*). Veterinary Parasitology, 153, 12–18. doi:10.1016/j.vetpar.2008.01.015

Milne, G., Fujimoto, C., Bean, T., Peters, H. J., Hemmington, M., Taylor, C., Fowkes, R. C., Martineau, H. M., Hamilton, C. M., Walker, M., Mitchell, J. A., Leger, E., Priestnall, S. L. & Webster, J. P. 2020. Infectious Causation of Abnormal Host Behavior: *Toxoplasma gondii* and Its Potential Association With Dopey Fox Syndrome. Frontiers in Psychiatry, 11. doi:10.3389/fpsyt.2020.513536

Montoya, J. G. & Liesenfeld, O. 2004. Toxoplasmosis. Lancet, 363, 1965–1976. doi:10.1016/s0140-6736(04)16412-x

Nobuto, K. 1966. Toxoplasmosis in Animal and Laboratory Diagnosis. Japan Agricultural Research Quarterly, 1, 11–18.

Prandovszky, E., Gaskell, E., Martin, H., Dubey, J. P., Webster, J. P. & Mcconkey, G. A. 2011. The Neurotropic Parasite *Toxoplasma gondii* Increases Dopamine Metabolism. Plos One, 6. doi:10.1371/journal.pone.0023866

R CORE TEAM 2022. R: A language and environment for statistical computing. R Foundation for Statistical Computing, Vienna, Austria. https://www.R-project.org/.

Sánchez-Sánchez, R., Vázquez, P., Ferre, I. & Ortega-Mora, L. M. 2018. Treatment of Toxoplasmosis and Neosporosis in Farm Ruminants: State of Knowledge and Future Trends. Current Topics in Medicinal Chemistry, 18, 1304–1323. doi:10.2174/1568026618666181002113617

Sarkunas, M., Velickaite, S., Bruzinskaite, R., Malakauskas, A. & Petkevicius, S. 2007. Faecal egg output and herbage contamination with infective larvae of species of *Ostertagia* and *Oesophagostomum* from naturally infected farmed sika deer *Cervus nippon* in Lithuania. Journal of Helminthology, 81, 79–84. doi:10.1017/s0022149x07241884

Skerrett, H. E. & Holland, C. V. 2001. Asymptomatic shedding of *Cryptosporidium* oocysts by red deer hinds and calves. Veterinary Parasitology, 94, 239–246. doi:10.1016/s0304-4017(00)00405-2

Tan, D. & Vyas, A. 2016. *Toxoplasma gondii* infection and testosterone congruently increase tolerance of male rats for risk of reward forfeiture. Hormones and Behavior, 79, 37–44. doi:10.1016/j.yhbeh.2016.01.003

Thiebaut, R., Leproust, S., Chene, G., Gilbert, R. & Grp, S. S. 2007. Effectiveness of prenatal treatment for congenital toxoplasmosis: a meta-analysis of individual patients’ data. Lancet, 369, 115–122.

Tong, W. H., Pavey, C., O’handley, R. & Vyas, A. 2021. Behavioral biology of *Toxoplasma gondii* infection. Parasites & Vectors, 14. doi:10.1186/s13071-020-04528-x

Torrey, E. F. & Yolken, R. H. 2013. Toxoplasma oocysts as a public health problem. Trends in Parasitology, 29, 380–384. doi:10.1016/j.pt.2013.06.001

Webster, J. P., Brunton, C. F. A. & Macdonald, D. W. 1994. Effect Of *Toxoplasma-gondii* Upon Neophobic Behavior In Wild Brown-Rats, *Rattus-Norvegicus*. Parasitology, 109, 37–43. doi:10.1017/s003118200007774x

Wilking, H., Thamm, M., Stark, K., Aebischer, T. & Seeber, F. 2016. Prevalence, incidence estimations, and risk factors of *Toxoplasma gondii* infection in Germany: a representative, cross-sectional, serological study. Scientific Reports, 6. doi:10.1038/srep22551

Williams, R., Morley, E., Hughes, J., Duncanson, P., Terry, R., Smith, J. & Hide, G. 2005 High levels of congenital transmission of *Toxoplasma gondii* in longitudinal and cross-sectional studies on sheep farms provides evidence of vertical transmission in ovine hosts. Parasitology, 130, 301–307. doi:10.1017/s0031182004006614

