## Supplementary material S1 for "Fallow deer approaching humans are also more likely to be seropositive for *Toxoplasma gondii*"

**Appendix 1**

The results of the validation test run to examine the relationships between S/P% values and sample dilution. The Sample ID is the code used to identify each sample. The dilution is the amount that the sample was diluted for the test (1:2 or 1:10). Each sample dilution was run on two different ELISA plates to give two runs (run 1, run 2). For each run we used the optical density given by the ELISA to calculate the S/P% values. We interpreted the results of each run separately using the thresholds given by the ELISA kit (<40 = Negative, >50 = Positive, 40-50 = Equivocal). We then compared the interpretation of each run, if the results agreed we classified the sample dilution positive or negative. If the results differed, we classified the sample dilution as equivocal.

|  |  | Run 1 |  | Run 2 |  | Overall |
| --- | --- | --- | --- | --- | --- | --- |
| Sample ID | Dilution | S/P % | Interpretation | S/P % | Interpretation | Interpretation |
| PB1 | 1:2 | 19.2251462 | Negative | 19.31608133 | Negative | Negative |
| PB1 | 1:10 | 7.090643275 | Negative | 6.74676525 | Negative | Negative |
| PB2 | 1:2 | 149.6345029 | Positive | 107.8558226 | Positive | Positive |
| PB2 | 1:10 | 174.6345029 | Positive | 90.4805915 | Positive | Positive |
| PB3 | 1:2 | 161.9152047 | Positive | 173.1053604 | Positive | Positive |
| PB3 | 1:10 | 130.9210526 | Positive | 114.6950092 | Positive | Positive |
| PB4 | 1:2 | 16.73976608 | Negative | 16.72828096 | Negative | Negative |
| PB4 | 1:10 | 5.921052632 | Negative | 5.637707948 | Negative | Negative |
| PB5 | 1:2 | 42.47076023 | Equivocal | 86.41404806 | Positive | Equivocal |
| PB5 | 1:10 | 72.5877193 | Positive | 107.8558226 | Positive | Positive |
| PB6 | 1:2 | 56.65204678 | Positive | 32.25508318 | Negative | Equivocal |
| PB6 | 1:10 | 39.54678363 | Negative | 29.11275416 | Negative | Negative |
| PB7 | 1:2 | 13.66959064 | Negative | 11.55268022 | Negative | Negative |
| PB7 | 1:10 | 14.25438596 | Negative | 10.44362292 | Negative | Negative |
| PB8 | 1:2 | 178.2894737 | Positive | 147.2273567 | Positive | Positive |
| PB8 | 1:10 | 132.2368421 | Positive | 99.53789279 | Positive | Positive |
| PB9 | 1:2 | 15.13157895 | Negative | 18.39186691 | Negative | Negative |
| PB9 | 1:10 | 7.529239766 | Negative | 0.092421442 | Negative | Negative |
| PB10 | 1:2 | 153.5818713 | Positive | 126.155268 | Positive | Positive |
| PB10 | 1:10 | 165.8625731 | Positive | 139.8336414 | Positive | Positive |

The vast majority of samples from the two dilutions come to the same classification. For two samples (PB5 and PB6) one of the dilutions was equivocal, however three out of the four times the sample was run it produced the same result. Therefore, we gathered enough support that motivated us to use dried blood collected on filter paper and eluted for our ELISA testing
