## Supplementary material S3 for "Fallow deer approaching humans are also more likely to be seropositive for *Toxoplasma gondii*"

**Appendix 3**

The effect plot depicting the effect of sex (a), serostatus (b), season (c), and year of study (d) on distance from buildings (y-axes, in meters) in deer monitored in the Phoenix Park, Dublin, as predicted by a linear mixed-effect model. The bars / shaded area represents the 95% marginal confidence intervals.

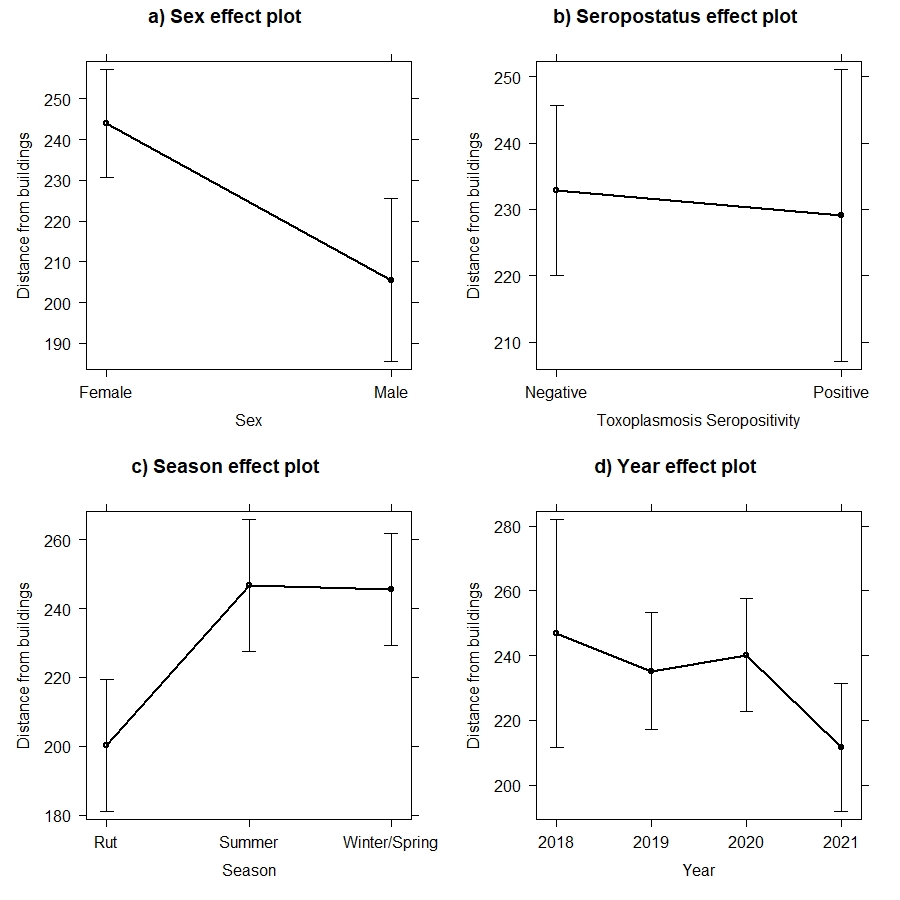
